# Gut microbiota-mediated alleviation of dextran sulfate sodium-induced colitis in mice

**DOI:** 10.1101/2023.08.07.552384

**Authors:** Eri Ikeda, Masaya Yamaguchi, Shigetada Kawabata

**Affiliations:** Department of Microbiology, Graduates School of Dentistry, Osaka University; Department of Molecular Immunology, Graduate School of Medical and Dental Sciences, Tokyo Medical and Dental University; Bioinformatics Research Unit, Graduates School of Dentistry, Osaka University; Center for Infectious Disease Education and Research (CiDER), Osaka University

## Abstract

**Summary:** We found a disease-resistant BALB/c substrain with lower susceptibility to DSS-induced colitis. Gut microbiota analysis using 16S rRNA analysis observed the expansion of butyrate-producing *Roseburia* species in mice with decreased susceptibility to the disease.

Gut dysbiosis characterized by an imbalanced microbiota is closed involved in the pathogenesis of a widespread gastrointestinal inflammatory disorder, inflammatory bowel disease. However, it is unclear how the complex intestinal microbiota affects resistance to mucosal inflammation. Here, we showed differences in the susceptibility of inbred BALB/c mice from three main distributors of laboratory animals to colitis; clinical symptoms, such as weight loss, disease activity index score, and colon shortening were assessed. Analysis of the gut microbiota using 16S rRNA sequencing revealed clear separation of the gut microbial composition among mice from the vendors. Notably, the abundance of the phylum *Actinobacteriota* was strongly associated with disease activity. We also observed the expansion of butyrate-producing *Roseburia* species in mice with decreased susceptibility to the disease. Further cohousing experiments showed that variation in clinical outcomes was more correlated with the gut microbiota than genetic variants among substrains from different suppliers. Targeting butyrate-producing bacteria could have therapeutic potential for ulcerative colitis.

## Introduction

Laboratory mice are important species for preclinical animal experiments in biomedical research and contribute to mechanistic studies and drug development in the context of various human diseases. Inbred mouse strains difference can be a reason of host immune characteristics as well as behavioural phenotypes(Bothe et al., 2005; Hensel et al., 2019). Numerous substrains have been derived from original inbred strains(Mekada et al., 2015). Substrains are defined as branches of an inbred strain produced by separated brother-sister mating over at least 20 generations from multiple vendors. During inbreeding, substrains from the classic inbred line diverge over time; thus, although these substrains are related, they are not identical because of housing conditions(Mahajan et al., 2016; Ussar et al., 2015; Warden et al., 2020).

The gastrointestinal tract of mammals is considered a primary reservoir for microorganisms that comprise vast and complex communities(Ley et al., 2008; Priya et al., 2022; Valles-Colomer et al., 2022). The gut microbiota, which includes hundreds of bacterial species, has been extensively investigated through epidemiological, physiological, and omics-based studies in humans(Baumgart and Carding, 2007; Fan and Pedersen, 2021; Glassner et al., 2020). Accumulating evidence suggests that disruption of the gut microbiome, which is known as dysbiosis, is closely connected to pathological states such as inflammation, autoimmune disorders, and cancer(Malesza et al., 2021; Yang et al., 2021; Zeng et al., 2017). Inflammatory bowel disease (IBD) is a multifactorial immune-mediated inflammatory disease that includes two conditions, Crohn’s disease and ulcerative colitis (Shah et al., 2023; Vich Vila et al., 2023). The gut dysbiosis associated with IBD is characterized by a lack of diversity, an increase in the levels of *Proteobacteria*, such as *Enterobacteriaceae* and *BIlophila*, and reduced levels of beneficial bacteria, such as short-chain fatty acid (SCFA)-producing *Clostridium*(Khan et al., 2022; Lee and Chang, 2021; Schirmer et al., 2018; Wright et al., 2015). SCFAs produced by gut bacteria act as coenzymes in fat and carbohydrate metabolism, thus exhibiting anti-inflammatory effects in IBD. Among the SCFAs, butyrate serves as a principal energy source for intestinal wound healing and barrier function. The gut microbiota of IBD patients exhibits a selective decrease in the levels of butyrate producers, and the colonocytes of IBD patients are incapable of transferring and utilizing butyrate(Wang et al., 2020).

The dextran sulfate sodium (DSS)-induced colitis model is routinely used as a principal mouse model of ulcerative colitis. DSS administration in drinking water mediates gut epithelial damage, which causes inflammation. Mouse strain and sex differences influence colitis susceptibility(Mahler et al., 1998). For example, similar to the clinical development of ulcerative colitis in humans, male mice are more likely to develop DSS-induced colitis than female mice(Chassaing et al., 2014; Goodman et al., 2020). Additionally, BALB/c mice require higher concentrations of DSS to induce colitis than C57BL/6J mice(Chassaing et al., 2014). C57BL/6 wild-type mice from the same inbred strain purchased from two vendors (substrains), Jackson laboratory and Taconic farms, showed different bacterial compositions, and Jackson mice showed significantly fewer species of bacteria(Ivanov et al., 2009). Colonization of the gut by a segmented filamentous bacterium was found only in Taconic mice, and the bacterium induced the production of inflammatory Th17 cells in the lamina propria of the small intestine, which resulted in increased resistance to *Citrobacter rodentium*-induced colitis(Ivanov et al., 2009). In addition, the composition of gut microbes that influence susceptibility to several diseases, including abdominal sepsis, vary among substrains(Hilbert et al., 2017).

Despite these advances in understanding, whether the gut microbiota is a primary factor determining disease susceptibility in DSS-induced colitis remains unknown. We compared mice from the same inbred strain (BALB/c) that were obtained from the three main distributors of laboratory animals in Japan and identified considerable variability in the presentation of DSS-induced colitis among mice from different vendors. We therefore quantified the influence of the faecal microbiota associated with mice from different vendors. We identified that mice from each vendor harboured a distinct gut microbiota. Thus, using a cohousing approach, we revealed that disease resistance mainly relies on the gut microbiota and not on genetic differences among substrains.

## Results

### The phenotype of DSS-induced colitis varied among mice from different vendors

We compared female six-to seven-week-old BALB/C wild-type mice obtained from three vendors: SLC, CLEA, and Charles River. Acute colitis was induced by the administration of 4% (w/v) DSS in drinking water for eight days. To investigate whether colitis induction by DSS was comparable among mice from different vendors, weight loss, rectal bleeding, stool consistency, colon shortening, and spleen enlargement were observed as clinical features of disease. Regarding weight loss, the body weights of mice from all vendors slightly increased during the first few days of the experiment. The weight of CLEA mice and Charles River mice gradually began to decrease afterward, and the body weights on day 8 were 14.4 ± 9.7 and 16.8 ± 5.7% below the initial weight, respectively (Fig. 1A). SLC mice showed little weight loss, and the body weight on day 8 was only 1.5 ± 4.1% below the initial weight. The level of weight loss in CLEA mice and Charles River mice significantly differed from that in SLC mice on day 7 (p = 0.04 and 0.03, respectively) and day 8 (p = 8.0×10^−3^and 4.0×10^−3^, respectively). Similarly, the DAI score, which comprises the degrees of weight loss and intestinal bleeding, was significantly higher (indicating more severe colitis) in CLEA and Charles River mice than in SLC mice on day 7 (p = 0.04 and 0.02, respectively) and day 8 (p = 0.02 and 0.02, respectively) (Fig. 1B). The disease activity index (DAI) score ranges from 0 to 12 (total score). Moreover, SLC mice did not exhibit considerable colon shortening (Fig. 1C, D) or spleen enlargement (Fig. 1E), unlike CLEA or Charles River mice. There was a trend of more colitis symptoms in Charles River mice than in CLEA mice, although significant differences in these symptoms of colitis were not observed. There were no excluded animals. Any unexpected adverse events haven’t seen in all animals.

**Figure 1.**
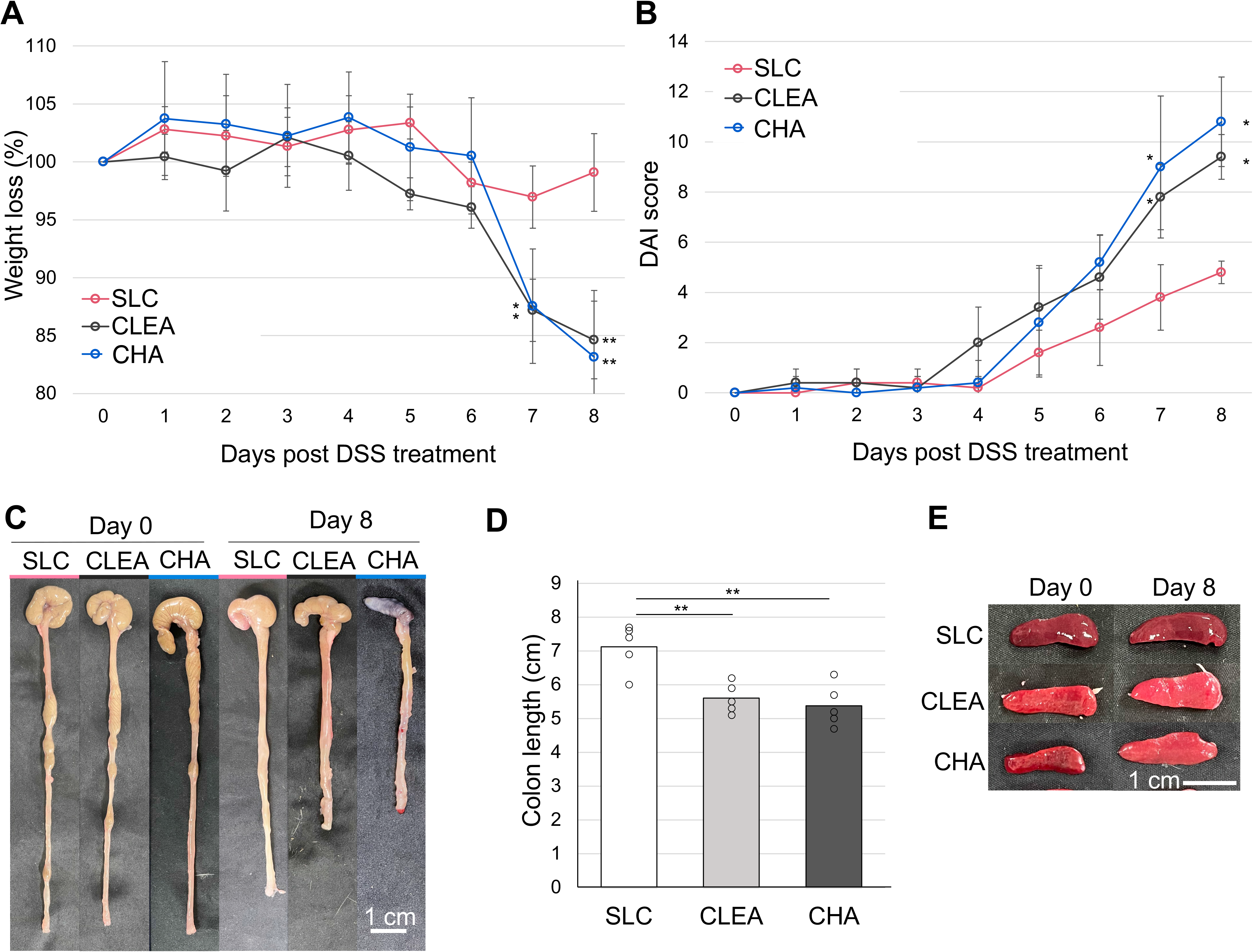
Clinical symptoms of colitis are highly variable among mice from different vendors. Mice from three commercial vendors, SLC, CLEA, and Charles River (CHA), were treated with 4% DSS for 8 days. Mice were evaluated daily, and weight loss and disease activity index scores were recorded. (A) Body weight changes, (n = 5). (B) DAI score, a score from 0 to 12. A higher number indicates more severe colitis (n = 5). (C) Gross images of the colon on days 0 (pre-DSS) and 8 (post-DSS). (D) Colon length was measured on day 8. (E) Gross images of the spleen on days 0 and 8. *P<0.05, **P<0.01.

### The gut microbial compositions of BALB/c mice largely varied among mice from different vendors

We compared the gut bacterial composition of mice from SLC, CLEA, and Charles River using 16S rRNA sequencing. Non-metric multidimensional scaling ordination using Horn-Morisita dissimilarities based on community membership indicated a clear separation of the microbiota among mice from the different vendors before DSS treatment (Fig. 2A). Consistent with previous studies, a significant decrease in microbial diversity, which is a characteristic of dysbiosis, was observed in CLEA and Charles River mice, whereas SLC mice did not show diversity changes after DSS administration (Fig. 2B). Microbial diversity was not significantly different among vendors either before or after DSS administration. Fifteen phyla were identified in total, as shown in Figure 2C. Contrary to our expectations, Charles River mice, but not SLC mice, were distinct from mice from other vendors before DSS treatment. In other words, SLC mice and CLEA mice were similar despite differences in disease severity. Notably, although *Bacteroidetes* and *Firmicutes* are two main phyla of the gut microbiota, the phylum *Firmicutes* was the most dominant in Charles River mice before DSS treatment, comprising up to 95%; hence, a low abundance of *Bacteroidetes* (2.2%) was observed. The phylum *Firmicutes* consists of the class *Bacilli* and class *Clostridia* (Supplemental Fig. 1), and the levels of *Clostridia* in mice before DSS treatment was significantly lower in mice from SLC compared to CLEA and Charles River (Fig. 2D), suggesting that the high *Firmicutes* abundance in Charles River mice was mainly due to the class *Clostridia*. The levels of *Clostridia* in mice after DSS treatment was not significantly different among vendors. Further investigation of the taxa comprising the class *Clostridia* was performed, and the levels of abundant families that were ≥ 1% abundant in at least one group are shown in Fig. 2E. A decrease in the levels of SCFA-producing *Clostridium* cluster IV (*Ruminococcaceae* and *Clostridia UCG-014*) and XIVa (*Lachnospiraceae*) is often associated with gut dysbiosis (Lopetuso et al., 2013). The levels of *Lachnospiraceae* and *Rumicococcaceae* in mice before DSS treatment were significantly lower in mice from SLC compared to CLEA and Charles River. The level of *Oscillospiraceae* in mice before DSS treatment was significantly lower in mice from SLC compared to Charles River. In addition to the presence of two main phyla, *Firmicutes* and *Bacteroidetes*, the analysis of the microbiota before DSS treatment showed a very strong association between the DAI inflammation score and *Actinobacteriota* proportion (R^2^ = 0.80), although the abundance of this microbe was quite low (Fig. 2F and Supplemental Fig. 2). To distinguish the bacterial taxa that are commonly found in mice from each vendor, differences in microbial taxa at the family level among vendors were calculated by the linear discriminant analysis (LDA) effect size (LEfSe) method as well as a heatmap (Fig. 3 and Supplemental Fig. 3). Eight taxa out of 13 that were significantly abundant in Charles River mice before DSS treatment were from the class *Clostridia* except *Lactobacillaceae*, *RF39, Deferribacteraceae*, *Erysipelatoclostridiaceae, Acholeplasmataceae*, and *Eggerthellaceae.* The family *Eggerthellaceae* is a member of phylum *Actinobacteriota,* and all *Actinobacteriota* bacteria found in this study belonged to the *Eggerthellaceae.* Contrary to the mice before DSS treatment, taxa comprising the class *Clostridia* were significantly lower levels in Charles Rimer mice after DSS treatment. Members of the family *Desulfovibrionaceae* and *Rikenllaceae* were significantly abundant in SLC mice both before and after DSS treatment. More detailed composition at genus level by LEfSe analysis indicated similar tendency (Fig. 4A and B). Members of the genus *Desulfovibrio* and *Bilophila,* both comprising family *Desulfovibrionaceae,* were significantly abundant in SLC mice both before and after DSS treatment. Twenty taxa out of 28 that were significantly abundant in Charles River mice before DSS treatment were from the class *Clostridia* and there were no abundant taxa in mice after DSS treatment. Furthermore, genus *Roseburia* is an only taxon comprising the class *Clastridia* in SLC mice after DSS treatment (Fig. 4B). Among the SCFAs producing bacteria, *Roseburia intestinalis* (a member of family *Lachnospiraceae*) and *Faecalibacterium prausnitzii* (a member of family *Oscillospiraceae*) are the primary butyrate producers in the human gut(Kasahara et al., 2018). The mean abundance of the genus *Roseburia* was higher in Charles River and CLEA mice than in SLC mice before DSS treatment, and the abundance increased in SLC mice after DSS treatment, whereas that in Charles River and CLEA mice decreased after DSS treatment. The relative abundance of the genus *Roseburia* was significantly higher in SLC mice compared to Charles River mice on day 8 (p = 0.04) (Fig 4 B and C). The genus *Faecalibacterium* was not identified in all samples.

**Figure 2.**
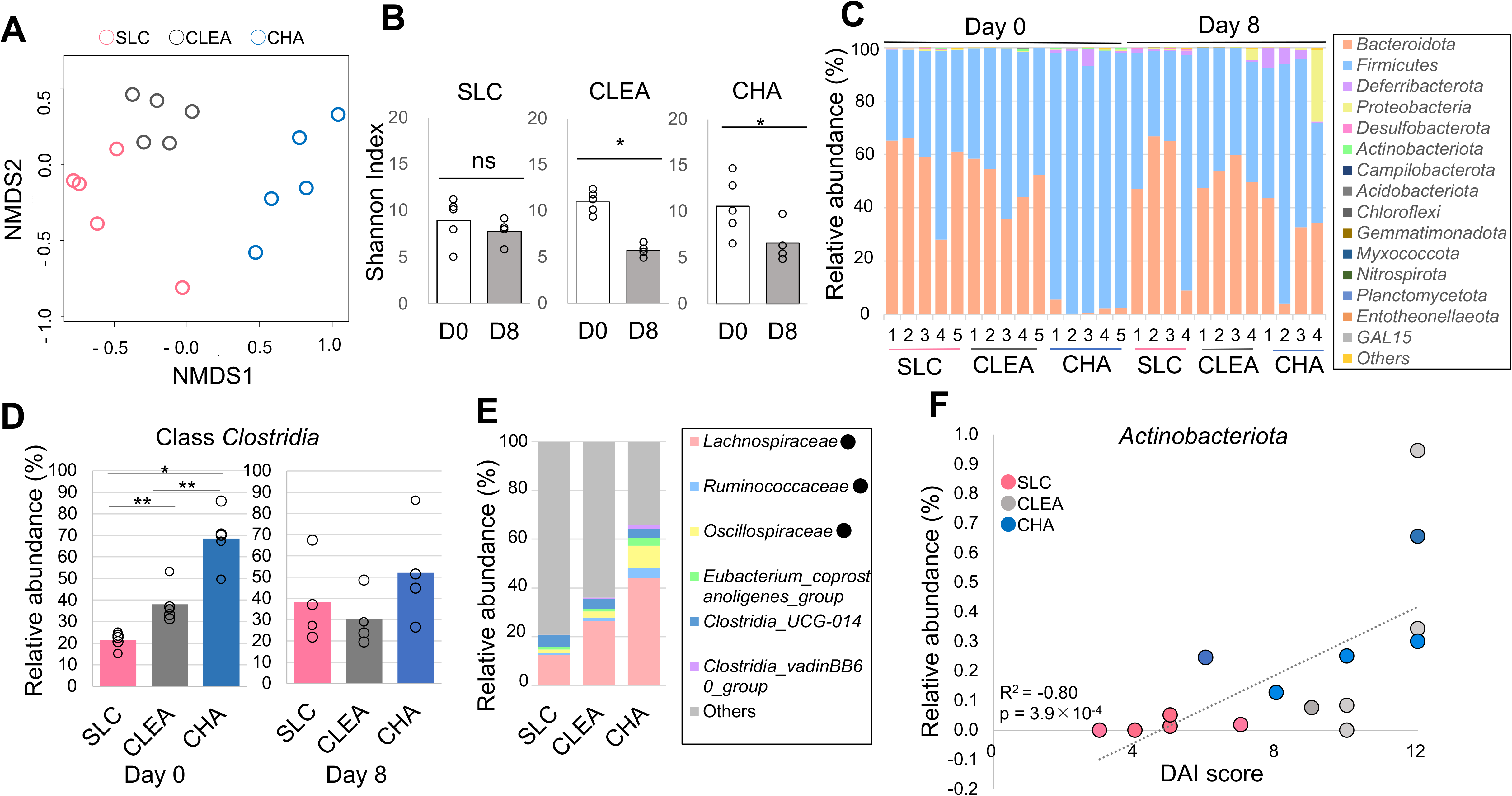
The gut microbiota in mice from three vendors assessed using 16S rRNA sequencing. The colorectal microbial composition in SLC, CLEA, and Charles River (CHA) mice treated with 4% DSS for 8 days was assessed using 16S rRNA amplicon sequencing. n = 4-5. (A) A nonmetric multidimensional scaling analysis identified a clear difference among mice vendor SLC (pink), CLEA (grey), and CHA (blue) before DSS treatment. (B) Dot plots show species-level microbial diversity measured by the Shannon diversity index. (C) Relative abundance of bacterial phyla presents in faeces on days 0 (pre-DSS) and 8 (post-DSS). (D) Relative abundance of class *Clostridia* comprising the phylum Firmicutes in faeces on days 0 (pre-DSS) and 8 (post-DSS). (E) Relative abundance of the bacterial family comprising the class *Clostridia* that was ≥ 1% abundant in at least one group of mice before DSS treatment. ●significantly different among vendors indicates. (F) Relationship between DAI score and relative abundance of phyla *Actinobacteriota* in mice from vendor SLC (pink) CLEA (grey), and CHA (blue) before DSS treatment (R^2^ = 0.83). *P<0.05, **P<0.01.

**Figure 3.**
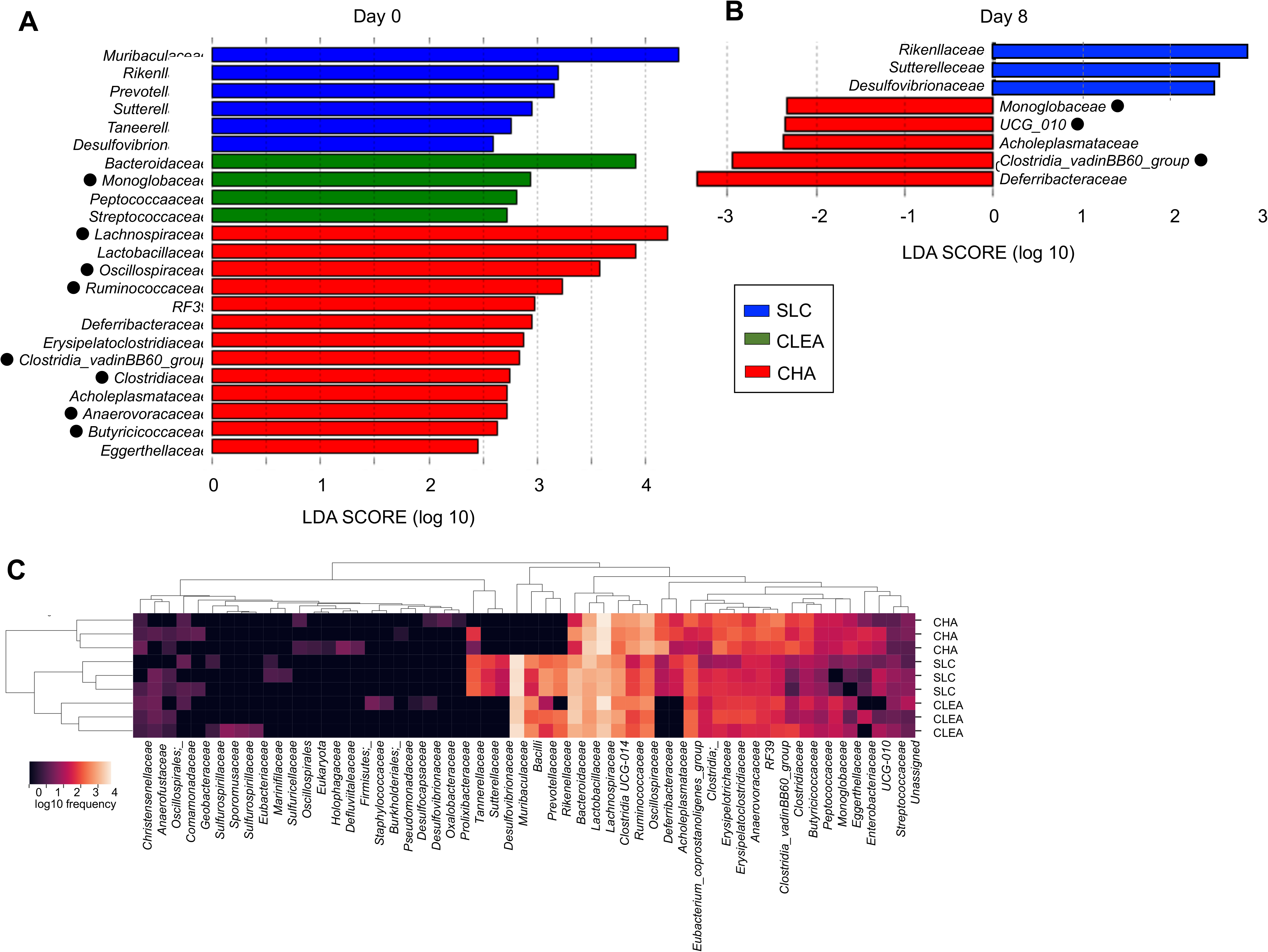
Differences in the microbiota of mice from the three vendors at the family level. Differences in microbiota taxa at the family level among mice from the three vendors were calculated by LDA effect size (LEfSe) on day 0 (A) and day 8 (B) (n=4-5). ● Bacterial taxa comprising the class *Clostridia*. (C) Heatmap showing bacterial family frequency distribution across mice from the three vendors before DSS treatment (n=3).

**Figure 4.**
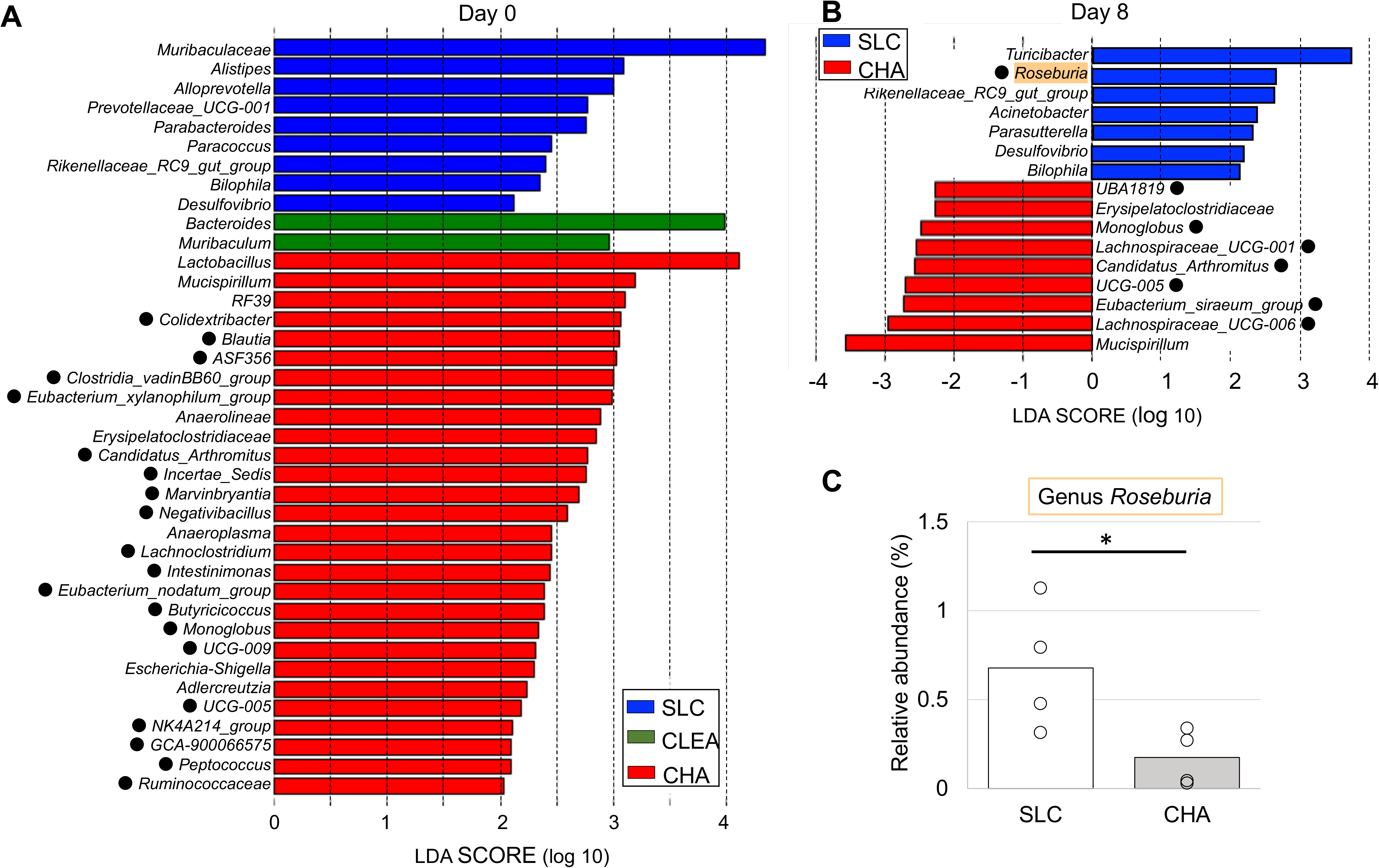
Differences in the microbiota of mice from the three vendors at the genus level. Differences in microbiota taxa at the genus level among mice from the three vendors were calculated by LDA effect size (LEfSe) on day 0 (A) and day 8 (B). ● Bacterial taxa comprising the class *Clostridia*. (C) Relative abundance of the butyrate-producing genus *Roseburia* of SLC and Charles River mice after DSS treatment (day 8).

### Commensal intestinal bacteria from mice with severe colitis influence the severity of disease in mice with milder colitis

Our gut microbiota analysis suggested that specific colonic microbes may induce colitis susceptibility. We next aimed to address the possibility that disease resistance to colitis in SLC mice might have been due to genetic variants among BALB/c mice substrains from the different vendors. Therefore, we cohoused SLC mice (mildest colitis symptoms) with Charles River mice (severest colitis symptoms) to allow horizontal bacterial transmission. SLC mice that were cohoused with Charles River mice for 4 weeks showed statistically higher DAI scores than SLC mice that were kept in separate cages (day 7, p = 0.03), suggesting that the gut microbiota, rather than the presence of genetic variants, is a dominant factor involved in the variable response to DSS (Fig. 5A). Regarding Charles River mice, although both Charles River mice housed separately and cohoused Charles River mice showed significantly higher DAI values than SLC mice (day 7, p = 0.03 and 0.01, respectively), no change was observed due to cohousing with SLC mice. Similarly, more colon shortening was observed in cohoused SLC mice, separately housed Charles River mice, and cohoused Charles River mice than in separately housed SLC mice (p = 1.6×10^−4^, 2.0×10^−3^, and 1.7×10^−3^, respectively) (Fig. 5B and C).

**Figure 5.**
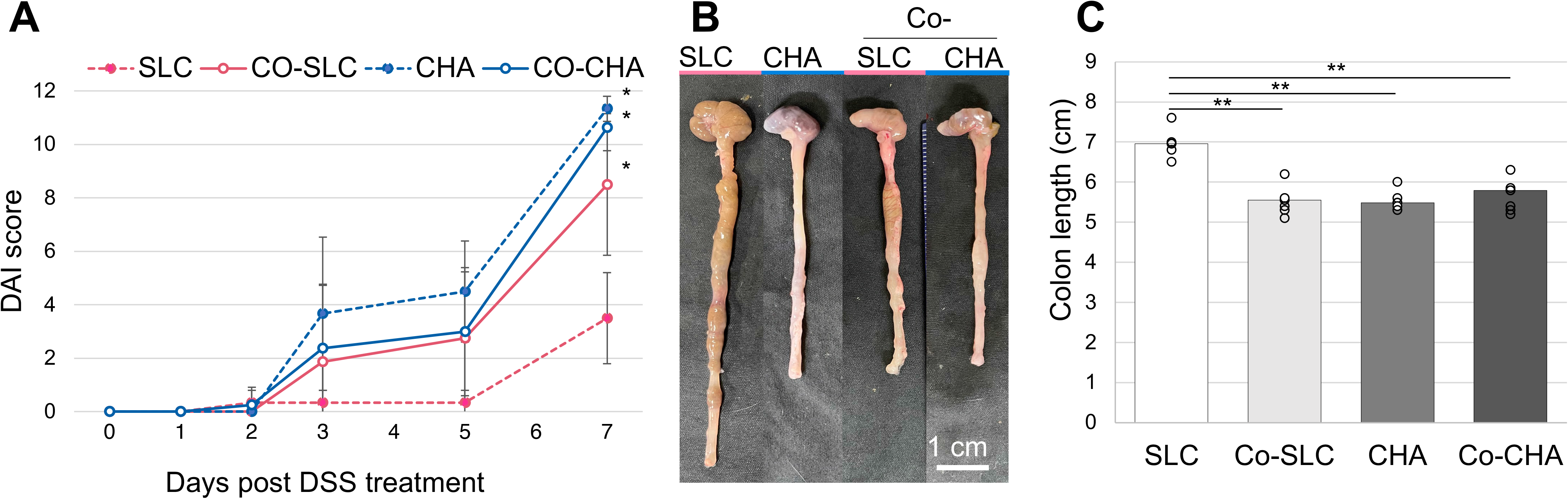
SLC mice cohoused with Charles River mice developed DSS-induced colitis. SLC mice (Co-SLC) and Charles River mice (Co-CHA) were cohoused for four weeks followed by 7 days of DSS administration (n=8). SLC mice (SLC) or Charles River mice (CHA) that were kept in separate cages were used as controls (n=6). Two independent experiments with identical results were combined. (A) DAI score. (B) Colon length was measured on day 7. *P<0.05, **P<0.01.

## Discussion

The DSS-induced colitis model is widely used because it can be established quickly and is simple(Chassaing et al., 2014). Here, we found clear differences in susceptibility to DSS-induced colitis among Japanese laboratory mice from three vendors. BALB/C mice purchased from SLC showed lower disease symptoms than mice from the other two vendors. The gut microbiota is known as an indispensable factor in gut inflammation(Shen et al., 2018). To quantify the connection between disease severity and the gut microbiota, the faeces of mice from three vendors collected before and after DSS treatment were compared using 16S rRNA sequencing, revealing that each group of mice from the different vendors harboured a distinct gut microbiota. Moreover, the cohousing data from the present study further show that the severity of DSS-induced colitis was mainly influenced by the gut microbiota.

The variability in DSS-induced colitis among individual mice from the same inbred strain has been documented in a large-scale animal experiment using genetically identical laboratory mice from a single animal facility(Forster et al., 2022). The researchers reported that the presence of specific gut bacteria was mainly responsible for the variable experimental outcomes in the DSS model. In humans, a large clinical cohort study examined genetic-microbial associations in healthy people who had different ancestral backgrounds but shared a relatively similar environment. The study showed that host genetics or ancestral backgrounds have a minor role in determining the gut microbiome; rather, the microbiota is shaped predominantly by lifestyle and is similar among individuals who share a relatively homogenous environment(Rothschild et al., 2018). IBD patients show a distinct distribution of certain bacterial taxa; IBD is accompanied by a decreased abundance of *Bacteroidetes, Firmicutes, Clostridia, Lactobacillus,* and *Ruminococcaceae* and an increased abundance of *Gammaproteobacteria* and *Enterobacteriaceae*(Gevers et al., 2014; Zheng et al., 2020). In particular, the *Firmicutes/Bacteroidetes* ratio is widely accepted to have vast influences on the maintenance of gut homeostasis, and an imbalance in these taxa can lead to various pathologies(Shen et al., 2018). Related to these epidemiological studies, our study revealed a higher percentage of *Firmicutes* in Charles River mice, and these bacteria were classified mainly into the class *Clostridia*. Among the *Clostridia*, members of *Clostridium* cluster IV (*Ruminococcaceae* and *Clostridia UCG-014*) and XIVa (*Lachnospiraceae*) are beneficial microbiota that produce SCFAs, including butyrate(Atarashi et al., 2013; Kabeerdoss et al., 2013; Vermeiren et al., 2012). In particular, *Roseburia intestinalis* is one of the primary butyrate producers in the human gut(Kasahara et al., 2018). The genus *Roseburia* consists of obligate gram-positive anaerobic bacteria, all of which are known to be SCFA producers(Tamanai-Shacoori et al., 2017). We reported that the abundance of the genus *Roseburia* increased precipitously in SLC mice after DSS treatment, while it was lower than that in mice from the other two vendors before DSS challenge. This implies that SCFA-producing *Roseburia* species may promote resistance to colitis in mice. Similarly, in a previous study that compared the gut microbiota of mice from two vendors (substrains) in the United States, colonization of the gut by colitis-resistant *Candidatus arthromitus* (a segmented filamentous bacterium and member of the family *Clostridiaceae*) was found in Taconic mice(Ivanov et al., 2009). The distribution of disease-resistant bacteria may not be ubiquitous but may have similar characteristics among different mice.

As previously discussed, we showed that few colitis symptoms were observed in SLC mice in our study. Although SLC is a major animal manufacturer, there have been few reports of studies using mice from SLC in the field of DSS-induced colitis research. One report using male C57BL/6 mice from SLC described body weight loss as well as a low DAI score, which supports our data(Tsuzuno et al., 2021). C57BL/6, BALB/c, and C3H/HeJ strains are known to be genetically susceptible to DSS-induced colitis(Eichele and Kharbanda, 2017). To our knowledge, this is the first report to describe a disease-resistant BALB/c substrain with lower disease susceptibility, and we showed that the susceptibility mainly relies on the gut microbiota. While our gut microbiota analysis using 16S rRNA analysis had limitations in the accurate characterization of species, the levels of the family *Muribaculaceae,* which was statistically abundant in SLC mice, have been reported to be negatively correlated with DSS-induced inflammation and further preserved in DSS-resistant *Dusp*6 knockout mice(Chang et al., 2021; Lagkouvardos et al., 2019; Shang et al., 2021). Considering these findings, members of the family *Muribaculaceae* could also be studied in the future as potential protective bacteria against disease.

In summary, a mouse substrain that was resistant to DSS-induced colitis was observed, and the severity of DSS-induced colitis was mainly influenced by the gut microbiota. DSS-induced colitis is one of the central preclinical models used in the gastrointestinal field. When studying disease susceptibility in wild-type and transgenic and/or genetically deficient mice, the mouse vendor and/or breeding conditions that influence the gut commensal microbiota may be the reason for variable outcomes. Moreover, our study supports the development of therapies targeting of butyrate or butyrate-producing bacteria for the prevention of ulcerative colitis.

## Materials and Methods

### Animals and DSS-induced colitis model

Female BALB/C wild-type mice were obtained from three animal vendors: SLC Japan (BALB/cCrSLC, Shizuoka, Japan), CLEA Japan (BALB/cAJcl, Tokyo, Japan), and Charles River Laboratories Japan (BALB/cAnNCrlCrlj Kanagawa, Japan). Mice were randomly assigned to cages before the experiment. Six-to seven-week-old mice were used except in the cohousing experiment. Acute colitis was induced by orally administrating 4% (w/v) DSS (molecular mass 36–50 kDa; MP biomedicals) in the drinking water for eight days. Body weight loss (0, none; 1, 1%–5%; 2, 5%–10%; 3, 10%–20%; 4, > 20%), rectal bleeding (0, normal; 2, haemoccult positive; 4, gross blood), and stool consistency (0, normal stool; 2, loose stool; 4, diarrhoea) were monitored daily. Mice These parameters were used to assess the colitis clinical score and the DAI. Mice lost more than 20% of their body weight during the experiment were euthanized. Mice were euthanized on day 8. Colon length was measured as the distance between the end of the caecum and proximal rectum. All animals that were used in this study were housed in groups of 3–6 mice, fed standard pellet diet, under a 12-hour light/dark cycle. Any research protocol was not registered before the study. This study does not involve human participants. All animal experiments were approved by the Institutional Animal Care and Use Committee of Tokyo Medical and Dental University (Protocol number: A2021-088C9) and the Animal Care and Use Committee of Osaka University Graduate School of Dentistry (R05-009-0). We confirm that all methods were carried out in accordance with relevant guidelines and regulations and that all methods are reported in accordance with the Animal Research: Reporting of In Vivo Experiments (ARRIVE) guidelines 2.0 for the reporting of animal experiments.

### 16S rRNA sequencing

Mouse faeces or colon luminal contents were collected on day 0 and day 8. Collected samples were stored at −80 °C until further use. Bacterial DNA was isolated using a NucleoSpin DNA stool kit (Takara Bio, Shiga, Japan). The V3-V4 regions of the 16S rRNA gene were amplified in each sample. Sequencing was performed on the Illumina MiSeq platform using a MiSeq Reagent Kit V3 (300 bp x 2) (Eurofins genomics, Tokyo, Japan).

Raw sequences were curated using the software package Qiime2. Sequences were assigned to operational taxonomic units using a cut-off = 0.03 and classified using the SILVA platform with a 70% confidence threshold. We used the LEfSe method(Segata et al., 2011) (http://huttenhower.sph.harvard.edu/lefse), which is used to perform a combined assessment of statistical significance and biological relevance.

### Cohousing

For the cohousing experiment, four-week-old BALB/C female mice purchased from SLC and Charles River were housed separately for one week in the same room and were fed the same diet before cohousing. Then, the mice were transferred into a new cage, and SLC mice and Charles River mice were cohoused for four weeks as described previously(Wirtz et al., 2017). SLC mice or Charles River mice that were kept in separate cages were used as controls. Acute colitis was induced with 4% (w/v) DSS for seven days afterwards.

### Statistics and reproducibility

Comparisons of two groups were performed using an unpaired *t* (parametric) test or a MannLWhitney *U* (nonparametric) test. Differences among more than three groups were evaluated using one-way analysis of variance for parametric analysis or the KruskalLWallis test for nonparametric analysis followed by Bonferroni correction (parametric) or Steel-Dwass correction (nonparametric). The normality of the data was analysed using the KolmogorovLSmirnov test. Homogeneity of variance was analysed using the F test (two groups) or Bartlett test (more than three groups). The sample distribution of the gut microbiota was analysed by a NMDS method. Correlation analysis was performed using the Pearson correlation coefficient. Error bars represent the standard deviation of a data set. All statistical analyses were performed with R statistical software.

**Data availability statement:** Metagenomic sequencing data were deposited in the DNA Data Bank of Japan (DDBJ) under project no. PRJDB15999.

## Acknowledgments

This work was supported by the Japan Society for the Promotion of Science KAKENHI under grants 20K18500 and 23K16015. The authors declare no competing financial interests.

## Abbreviations

DAI: disease activity index
DSS: dextran sulfate sodium
IBD: inflammatory bowel disease
LDA: linear discriminant analysis
LEfSe: linear discriminant analysis effect size
SCFA: short-chain fatty acid.

## Notes

### Competing Interest Statement

The authors have declared no competing interest.

## References

Atarashi, K., T. Tanoue, K. Oshima, W. Suda, Y. Nagano, H. Nishikawa, S. Fukuda, T. Saito, S. Narushima, K. Hase, S. Kim, J.V. Fritz, P. Wilmes, S. Ueha, K. Matsushima, H. Ohno, B. Olle, S. Sakaguchi, T. Taniguchi, H. Morita, M. Hattori, and K. Honda. 2013. Treg induction by a rationally selected mixture of Clostridia strains from the human microbiota. Nature 500:232–236.

Baumgart, D.C., and S.R. Carding. 2007. Inflammatory bowel disease: cause and immunobiology. Lancet 369:1627–1640.

Bothe, G.W., V.J. Bolivar, M.J. Vedder, and J.G. Geistfeld. 2005. Behavioral differences among fourteen inbred mouse strains commonly used as disease models. Comp Med 55:326–334.

Chang, C.S., Y.C. Liao, C.T. Huang, C.M. Lin, C.H.Y. Cheung, J.W. Ruan, W.H. Yu, Y.T. Tsai, I.J. Lin, C.H. Huang, J.S. Liou, Y.H. Chou, H.J. Chien, H.L. Chuang, H.F. Juan, H.C. Huang, H.L. Chan, Y.C. Liao, S.C. Tang, Y.W. Su, T.H. Tan, A.J. Baumler, and C.Y. Kao. 2021. Identification of a gut microbiota member that ameliorates DSS-induced colitis in intestinal barrier enhanced Dusp6-deficient mice. Cell Rep 37:110016.

Chassaing, B., J.D. Aitken, M. Malleshappa, and M. Vijay-Kumar. 2014. Dextran sulfate sodium (DSS)-induced colitis in mice. Curr Protoc Immunol 104:15 251-15 25 14.

Eichele, D.D., and K.K. Kharbanda. 2017. Dextran sodium sulfate colitis murine model: An indispensable tool for advancing our understanding of inflammatory bowel diseases pathogenesis. World J Gastroenterol 23:6016–6029.

Fan, Y., and O. Pedersen. 2021. Gut microbiota in human metabolic health and disease. Nat Rev Microbiol 19:55–71.

Forster, S.C., S. Clare, B.S. Beresford-Jones, K. Harcourt, G. Notley, M.D. Stares, N. Kumar, A.T. Soderholm, A. Adoum, H. Wong, B. Moron, C. Brandt, G. Dougan, D.J. Adams, K.J. Maloy, V.A. Pedicord, and T.D. Lawley. 2022. Identification of gut microbial species linked with disease variability in a widely used mouse model of colitis. Nat Microbiol 7:590–599.

Gevers, D., S. Kugathasan, L.A. Denson, Y. Vazquez-Baeza, W. Van Treuren, B. Ren, E. Schwager, D. Knights, S.J. Song, M. Yassour, X.C. Morgan, A.D. Kostic, C. Luo, A. Gonzalez, D. McDonald, Y. Haberman, T. Walters, S. Baker, J. Rosh, M. Stephens, M. Heyman, J. Markowitz, R. Baldassano, A. Griffiths, F. Sylvester, D. Mack, S. Kim, W. Crandall, J. Hyams, C. Huttenhower, R. Knight, and R.J. Xavier. 2014. The treatment-naive microbiome in new-onset Crohn’s disease. Cell Host Microbe 15:382–392.

Glassner, K.L., B.P. Abraham, and E.M.M. Quigley. 2020. The microbiome and inflammatory bowel disease. J Allergy Clin Immunol 145:16–27.

Goodman, W.A., I.P. Erkkila, and T.T. Pizarro. 2020. Sex matters: impact on pathogenesis, presentation and treatment of inflammatory bowel disease. Nat Rev Gastroenterol Hepatol 17:740–754.

Hensel, J.A., V. Khattar, R. Ashton, and S. Ponnazhagan. 2019. Characterization of immune cell subtypes in three commonly used mouse strains reveals gender and strain-specific variations. Lab Invest 99:93–106.

Hilbert, T., F. Steinhagen, S. Senzig, N. Cramer, I. Bekeredjian-Ding, M. Parcina, G. Baumgarten, A. Hoeft, S. Frede, O. Boehm, and S. Klaschik. 2017. Vendor effects on murine gut microbiota influence experimental abdominal sepsis. J Surg Res 211:126–136.

Ivanov, II, K. Atarashi, N. Manel, E.L. Brodie, T. Shima, U. Karaoz, D. Wei, K.C. Goldfarb, C.A. Santee, S.V. Lynch, T. Tanoue, A. Imaoka, K. Itoh, K. Takeda, Y. Umesaki, K. Honda, and D.R. Littman. 2009. Induction of intestinal Th17 cells by segmented filamentous bacteria. Cell 139:485–498.

Kabeerdoss, J., V. Sankaran, S. Pugazhendhi, and B.S. Ramakrishna. 2013. Clostridium leptum group bacteria abundance and diversity in the fecal microbiota of patients with inflammatory bowel disease: a case-control study in India. BMC Gastroenterol 13:20.

Kasahara, K., K.A. Krautkramer, E. Org, K.A. Romano, R.L. Kerby, E.I. Vivas, M. Mehrabian, J.M. Denu, F. Backhed, A.J. Lusis, and F.E. Rey. 2018. Interactions between Roseburia intestinalis and diet modulate atherogenesis in a murine model. Nat Microbiol 3:1461-1471.

Khan, I., J. Wei, A. Li, Z. Liu, P. Yang, Y. Jing, X. Chen, T. Zhao, Y. Bai, L. Zha, C. Li, N. Ullah, T. Che, and C. Zhang. 2022. Lactobacillus plantarum strains attenuated DSS-induced colitis in mice by modulating the gut microbiota and immune response. Int Microbiol 25:587–603.

Lagkouvardos, I., T.R. Lesker, T.C.A. Hitch, E.J.C. Galvez, N. Smit, K. Neuhaus, J. Wang, J.F. Baines, B. Abt, B. Stecher, J. Overmann, T. Strowig, and T. Clavel. 2019. Sequence and cultivation study of Muribaculaceae reveals novel species, host preference, and functional potential of this yet undescribed family. Microbiome 7:28.

Lee, M., and E.B. Chang. 2021. Inflammatory Bowel Diseases (IBD) and the Microbiome-Searching the Crime Scene for Clues. Gastroenterology 160:524–537.

Ley, R.E., M. Hamady, C. Lozupone, P.J. Turnbaugh, R.R. Ramey, J.S. Bircher, M.L. Schlegel, T.A. Tucker, M.D. Schrenzel, R. Knight, and J.I. Gordon. 2008. Evolution of mammals and their gut microbes. Science 320:1647–1651.

Lopetuso, L.R., F. Scaldaferri, V. Petito, and A. Gasbarrini. 2013. Commensal Clostridia: leading players in the maintenance of gut homeostasis. Gut Pathog 5:23.

Mahajan, V.S., E. Demissie, H. Mattoo, V. Viswanadham, A. Varki, R. Morris, and S. Pillai. 2016. Striking Immune Phenotypes in Gene-Targeted Mice Are Driven by a Copy-Number Variant Originating from a Commercially Available C57BL/6 Strain. Cell Rep 15:1901–1909.

Mahler, M., I.J. Bristol, E.H. Leiter, A.E. Workman, E.H. Birkenmeier, C.O. Elson, and J.P. Sundberg. 1998. Differential susceptibility of inbred mouse strains to dextran sulfate sodium-induced colitis. Am J Physiol 274:G544–551.

Malesza, I.J., M. Malesza, J. Walkowiak, N. Mussin, D. Walkowiak, R. Aringazina, J. Bartkowiak-Wieczorek, and E. Madry. 2021. High-Fat, Western-Style Diet, Systemic Inflammation, and Gut Microbiota: A Narrative Review. Cells 10:

Mekada, K., M. Hirose, A. Murakami, and A. Yoshiki. 2015. Development of SNP markers for C57BL/6N-derived mouse inbred strains. Exp Anim 64:91–100.

Priya, S., M.B. Burns, T. Ward, R.A.T. Mars, B. Adamowicz, E.F. Lock, P.C. Kashyap, D. Knights, and R. Blekhman. 2022. Identification of shared and disease-specific host gene-microbiome associations across human diseases using multi-omic integration. Nat Microbiol 7:780–795.

Rothschild, D., O. Weissbrod, E. Barkan, A. Kurilshikov, T. Korem, D. Zeevi, P.I. Costea, A. Godneva, I.N. Kalka, N. Bar, S. Shilo, D. Lador, A.V. Vila, N. Zmora, M. Pevsner-Fischer, D. Israeli, N. Kosower, G. Malka, B.C. Wolf, T. Avnit-Sagi, M. Lotan-Pompan, A. Weinberger, Z. Halpern, S. Carmi, J. Fu, C. Wijmenga, A. Zhernakova, E. Elinav, and E. Segal. 2018. Environment dominates over host genetics in shaping human gut microbiota. Nature 555:210–215.

Schirmer, M., E.A. Franzosa, J. Lloyd-Price, L.J. McIver, R. Schwager, T.W. Poon, A.N. Ananthakrishnan, E. Andrews, G. Barron, K. Lake, M. Prasad, J. Sauk, B. Stevens, R.G. Wilson, J. Braun, L.A. Denson, S. Kugathasan, D.P.B. McGovern, H. Vlamakis, R.J. Xavier, and C. Huttenhower. 2018. Dynamics of metatranscription in the inflammatory bowel disease gut microbiome. Nat Microbiol 3:337–346.

Segata, N., J. Izard, L. Waldron, D. Gevers, L. Miropolsky, W.S. Garrett, and C. Huttenhower. 2011. Metagenomic biomarker discovery and explanation. Genome Biol 12:R60.

Shah, S.A., L. Deng, J. Thorsen, A.G. Pedersen, M.B. Dion, J.L. Castro-Mejia, R. Silins, F.O. Romme, R. Sausset, L.E. Jessen, E.O. Ndela, M. Hjelmso, M.A. Rasmussen, T.A. Redgwell, C. Leal Rodriguez, G. Vestergaard, Y. Zhang, B. Chawes, K. Bonnelykke, S.J. Sorensen, H. Bisgaard, F. Enault, J. Stokholm, S. Moineau, M.A. Petit, and D.S. Nielsen. 2023. Expanding known viral diversity in the healthy infant gut. Nat Microbiol 8:986–998.

Shang, L., H. Liu, H. Yu, M. Chen, T. Yang, X. Zeng, and S. Qiao. 2021. Core Altered Microorganisms in Colitis Mouse Model: A Comprehensive Time-Point and Fecal Microbiota Transplantation Analysis. Antibiotics (Basel) 10:

Shen, Z.H., C.X. Zhu, Y.S. Quan, Z.Y. Yang, S. Wu, W.W. Luo, B. Tan, and X.Y. Wang. 2018. Relationship between intestinal microbiota and ulcerative colitis: Mechanisms and clinical application of probiotics and fecal microbiota transplantation. World J Gastroenterol 24:5–14.

Tamanai-Shacoori, Z., I. Smida, L. Bousarghin, O. Loreal, V. Meuric, S.B. Fong, M. Bonnaure-Mallet, and A. Jolivet-Gougeon. 2017. Roseburia spp.: a marker of health? Future Microbiol 12:157–170.

Tsuzuno, T., N. Takahashi, M. Yamada-Hara, M. Yokoji-Takeuchi, B. Sulijaya, Y. Aoki-Nonaka, A. Matsugishi, K. Katakura, K. Tabeta, and K. Yamazaki. 2021. Ingestion of Porphyromonas gingivalis exacerbates colitis via intestinal epithelial barrier disruption in mice. J Periodontal Res 56:275–288.

Ussar, S., N.W. Griffin, O. Bezy, S. Fujisaka, S. Vienberg, S. Softic, L. Deng, L. Bry, J.I. Gordon, and C.R. Kahn. 2015. Interactions between Gut Microbiota, Host Genetics and Diet Modulate the Predisposition to Obesity and Metabolic Syndrome. Cell Metab 22:516–530.

Valles-Colomer, M., R. Bacigalupe, S. Vieira-Silva, S. Suzuki, Y. Darzi, R.Y. Tito, T. Yamada, N. Segata, J. Raes, and G. Falony. 2022. Variation and transmission of the human gut microbiota across multiple familial generations. Nat Microbiol 7:87–96.

Vermeiren, J., P. Van den Abbeele, D. Laukens, L.K. Vigsnaes, M. De Vos, N. Boon, and T. Van de Wiele. 2012. Decreased colonization of fecal Clostridium coccoides/Eubacterium rectale species from ulcerative colitis patients in an in vitro dynamic gut model with mucin environment. FEMS Microbiol Ecol 79:685–696.

Vich Vila, A., S. Hu, S. Andreu-Sanchez, V. Collij, B.H. Jansen, H.E. Augustijn, L.A. Bolte, R. Ruigrok, G. Abu-Ali, C. Giallourakis, J. Schneider, J. Parkinson, A. Al-Garawi, A. Zhernakova, R. Gacesa, J. Fu, and R.K. Weersma. 2023. Faecal metabolome and its determinants in inflammatory bowel disease. Gut

Wang, R.X., J.S. Lee, E.L. Campbell, and S.P. Colgan. 2020. Microbiota-derived butyrate dynamically regulates intestinal homeostasis through regulation of actin-associated protein synaptopodin. Proc Natl Acad Sci U S A 117:11648-11657.

Warden, A.S., A. DaCosta, S. Mason, Y.A. Blednov, R.D. Mayfield, and R.A. Harris. 2020. Inbred Substrain Differences Influence Neuroimmune Response and Drinking Behavior. Alcohol Clin Exp Res 44:1760–1768.

Wirtz, S., V. Popp, M. Kindermann, K. Gerlach, B. Weigmann, S. Fichtner-Feigl, and M.F. Neurath. 2017. Chemically induced mouse models of acute and chronic intestinal inflammation. Nat Protoc 12:1295–1309.

Wright, E.K., M.A. Kamm, S.M. Teo, M. Inouye, J. Wagner, and C.D. Kirkwood. 2015. Recent advances in characterizing the gastrointestinal microbiome in Crohn’s disease: a systematic review. Inflamm Bowel Dis 21:1219–1228.

Yang, Y., L. Du, D. Shi, C. Kong, J. Liu, G. Liu, X. Li, and Y. Ma. 2021. Dysbiosis of human gut microbiome in young-onset colorectal cancer. Nat Commun 12:6757.

Zeng, M.Y., N. Inohara, and G. Nunez. 2017. Mechanisms of inflammation-driven bacterial dysbiosis in the gut. Mucosal Immunol 10:18–26.

Zheng, D., T. Liwinski, and E. Elinav. 2020. Interaction between microbiota and immunity in health and disease. Cell Res 30:492–506.

