## supplementaly file for "Gut microbiota-mediated alleviation of dextran sulfate sodium-induced colitis in mice"

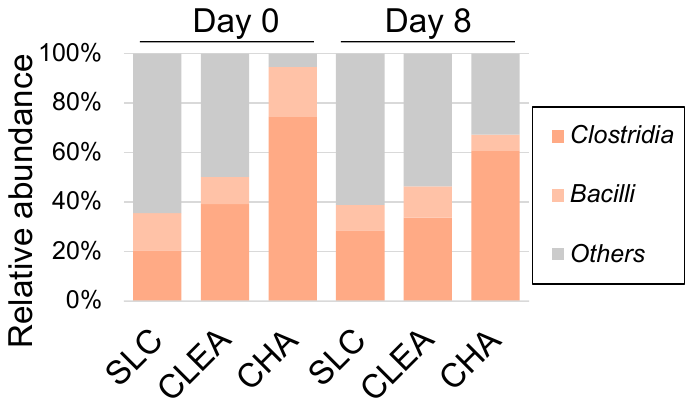

**Supplemental Figure 1 Relative abundance of bacterial classes comprising the phylum Firmicutes.**

Microbial classes comprising the phylum Firmicutes in faeces on days 0 (pre-DSS) and 8 (post-DSS).

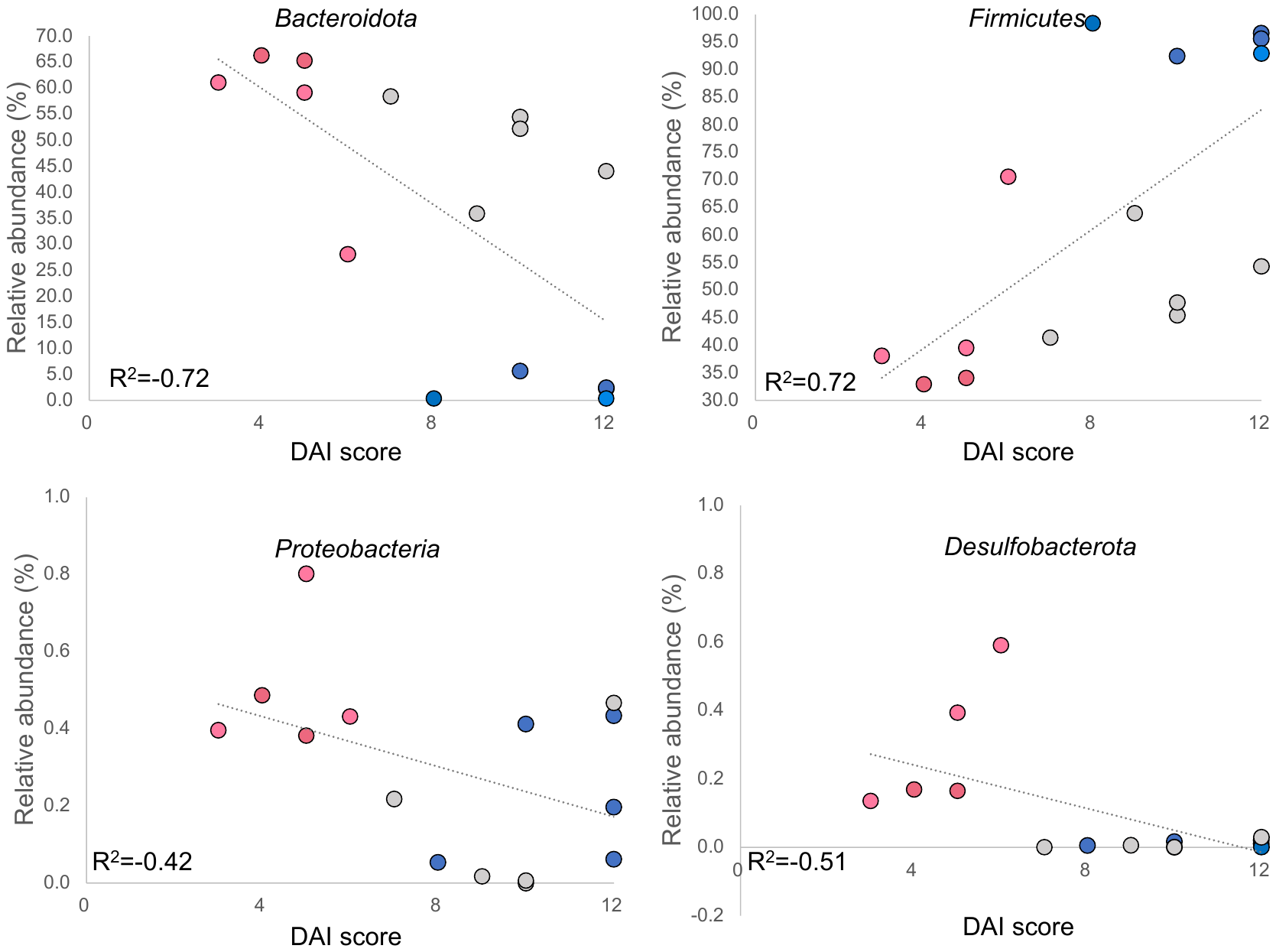

**Supplemental Figure 2 Relationship between DAI score and relative abundance of each phylum in mice before DSS treatment.** Dots indicate mice vendor SLC (pink) CLEA (grey), and CHA (blue). The Phyla that were present in all group were used. (A) Bacteroidota, (B) Firmicutes, (c)Proteobacteria, and (D) Desulfobateria.

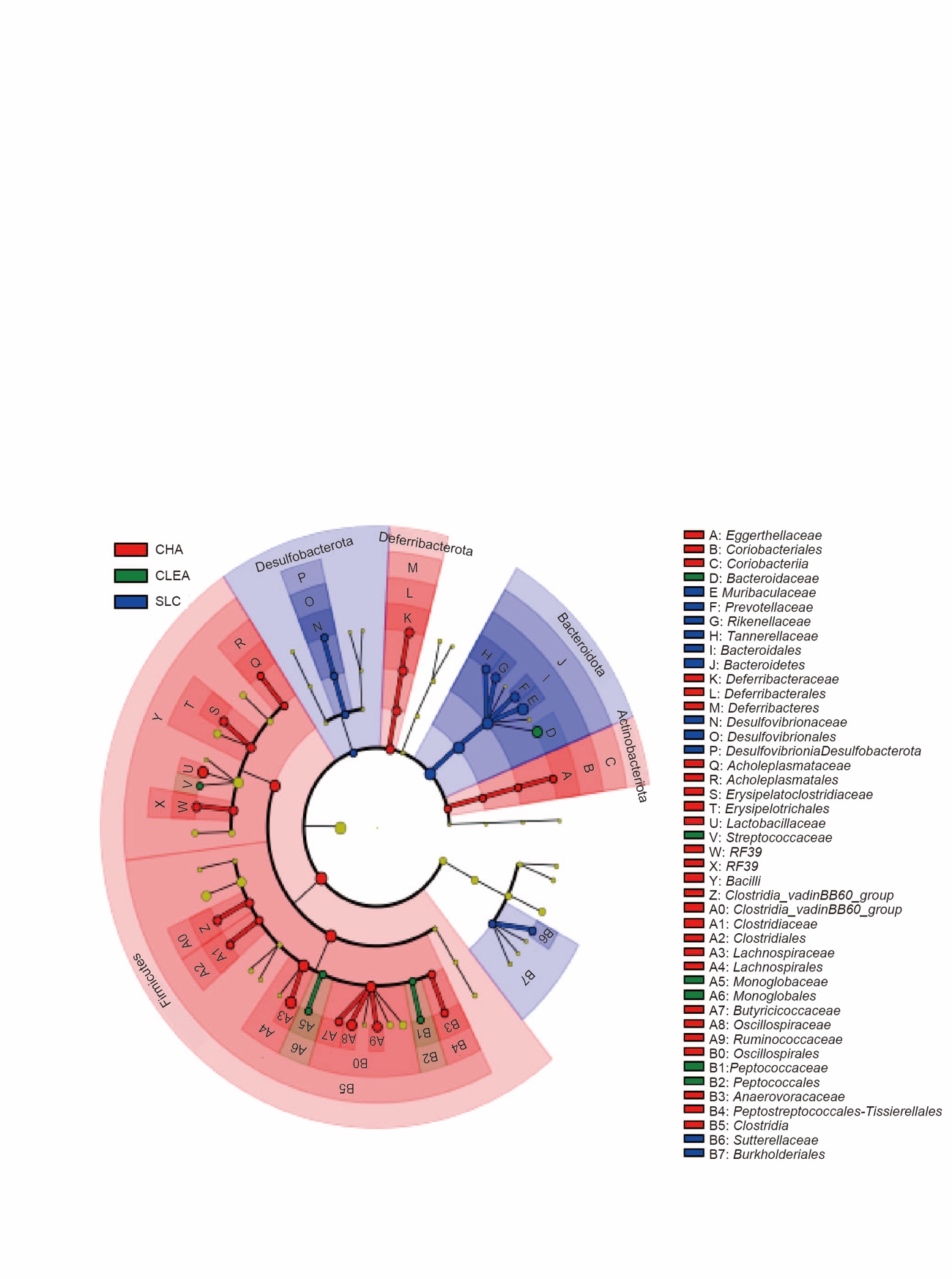

**Supplemental Figure 3 Taxonomic cladogram for LEfSe analysis of gut microbiota.** LEfSe cladogram demonstrating taxonomic differences among the vendors on day 0. Taxa and nodes highlighted in red, green, and blue were significantly more abundant in Charles River (CHA), CLEA, and SLC mice respectively. The diameter of each circle is proportional to the abundance of the taxon. Only differentially abundant taxa at the family or higher taxonomic ranks were indicated. Nodes remaining yellow indicate taxa that were not significantly differentially represented.
